# Synthetic cassettes for pH-mediated sensing, counting and containment

**DOI:** 10.1101/740902

**Authors:** Finn Stirling, Alexander Naydich, Juliet Bramante, Rachel Barocio, Michael Certo, Hannah Wellington, Elizabeth Redfield, Samuel O’Keefe, Sherry Gao, Adam Cusolito, Jeffrey Way, Pamela Silver

## Abstract

As pH is fundamental to all biological processes, pH-responsive bacterial genetic circuits enable precise sensing in any environment. Where unintentional release of engineered bacteria poses a concern, coupling pH sensing to expression of a toxin creates an effective bacterial containment system. Here, we present a pH-sensitive kill switch (acidic Termination of Replicating Population; acidTRP), based on the *E. coli asr* promoter, with a survival ratio of less than 1 in 10^6^. We integrate acidTRP with cryodeath to produce a two-factor containment system with a combined survival ratio of less than 1 in 10^11^ whilst maintaining evolutionary stability. We further develop a pulse-counting circuit with single cell readout for each administered stimulus pulse. We use this pulse-counter to record multiple pH changes and combine it with acidTRP to make a two-count acid-sensitive kill switch. These results demonstrate the ability to build complex genetic systems for biological containment.

## Introduction

pH is an essential aspect of all biological processes. Sensing and responding to pH is therefore a powerful tool in the arsenal of a synthetic biologist, allowing for an engineered strain to be controlled by a variety of environments. For example, bacterial species that travel through the mammalian digestive system are exposed to multiple levels of acidity (Fallinborf, 1999; McConnell et al., 2007). Endocytosis of pathogens by the human immune system results in a decrease in environmental pH for the pathogenic bacteria (Geisow & Evans, 1984; Wang et al., 2017). The soil microbiome senses and responds to the fluctuations in pH (Rousk et al., 2010).

*E. coli* responds to a wide array of stressful situations, allowing it to thrive in a variety of conditions (Arsène et al., 2000; Babai & Ron, 1998; Baranyi et al., 2014; Bläsi & Young, 1996; Rodrigues & Rodrigues, 2018). In response to a rapid reduction of pH, the acid shock response gene *asr* is upregulated (Motieju, 2009; Šeputiene et al., 2004). The P*_asr_* promoter is pH-responsive allowing pH-dependent control of the expression of a gene of interest (Hoynes-O’Connor et al., 2017).

A key aspect of bacterial control is the capacity to dictate what environment an engineered strain is capable of surviving in. For engineered bacterial strains to be deployed in clinical or environmental settings outside of a research laboratory, there must be a degree of confidence that they cannot escape to surrounding environments. Previously engineered forms of containment have used kill switches that rely on a small molecule survival factor to be provided by human intervention to ensure survival by i) allowing for expression of an essential gene (Cai et al., 2015); ii) regulation of a toxic gene (Caliando & Voigt, 2015; Contreras et al., 1991; Dirks et al., 2007; Ronchel & Ramos, 2001); iii) supplementation of an auxotrophy for a non-natural amino acid (Rovner et al., 2015), or a combination of multiple methods (Chan et al., 2015; Gallagher et al., 2015). Coupling containment to an environmental condition such as pH is an alternative way of controlling bacterial survival.

For any kill switch, the evolutionary stability of the system is paramount to its effectiveness. Often, a containment system designed to control bacterial growth causes a fitness defect, and hence a mutant defective in the killing function would have a growth advantage (Chan et al., 2015). If cell death is initiated by the up-regulation of a toxic protein, instability of a kill switch can be attributed to the basal expression of the toxic gene even in permissive conditions. Previously we and others have shown that the inclusion of an antitoxin, and careful balancing of toxin and antitoxin expression levels across relevant conditions, can mitigate this effect, resulting in a kill switch that is evolutionarily stable over biologically relevant time periods (Gallagher et al., 2015; Stirling et al., 2017).

A containment system must also have a survival ratio (defined as the fraction of the population able to survive in the non-permissive environment) low enough that the probability of an escape event is essentially zero. Previously published kill switches that respond to intrinsic characteristics of their environment have had survival ratios no lower than 10^-6^ (Piraner et al., 2016; Stirling et al., 2017), corresponding roughly to the mutation rate of a single gene (Lee et al., 2012). In practical environmental applications, population sizes are much greater than 10^6^ cells, so even lower survival ratios are desirable. Here, we present a containment system with a survival ratio of less than 1 in 10^11^.

A pulse counter is a specific type of recording circuit which tracks the number of discrete pulses of a single stimulus. Biological pulse counters might be used to monitor extracellular factors, such as fluctuations in antibiotic concentration at remote infection sites, or the rate of intracellular processes, such as bacterial division or metabolism. They may also be used to program a cellular response after a specific number of exposures. An effective cellular pulse counter demands three key characteristics: i) the circuit must exhibit a high signal-to-noise ratio, preventing advancement of the counter in the absence of a stimulus; ii) the circuit must produce a clear digital readout, such that the count can be interpreted on a single-cell basis; and iii) the circuit must advance the count only once for each discrete stimulus pulse, regardless of how long the pulse lasts. This last feature has been especially challenging for previous synthetic biological counters, since in the absence of negative feedback, multiple counts may be recorded if the duration and concentration of the stimulus pulse are not carefully controlled (Friedland et al., 2009). Since the first attempts at constructing pulse counter circuits (Friedland et al., 2009), several mechanisms have been proposed for how a counter might avoid erroneously registering multiple counts for a single stimulus by delaying its response until the falling edge of a stimulus pulse, but none have been implemented to date (Noman et al., 2016; Subsoontorn & Endy, 2012). Here, we design a robust pulse counting circuit and combine it with our kill switch to demonstrate our ability to program cellular behavior in response to a specific number of recurrences of a stimulus.

## Results

### Construction and Testing of the pH sensitive kill switch acidTRP

We constructed a pH-sensitive kill switch, Acid Termination of Replicating Population (or AcidTRP), based on the *E. coli* P*_asr_* promoter. P*_asr_* is a pH-sensitive promoter native to *E. coli* that is repressed at pH 7 and induced at pH 5 (Figure 1A). It controls expression of *asr* (acid shock RNA), encoding a protein of unknown function that is heavily upregulated during the acid shock response (Šeputiene et al., 2004). Regulation of P*_asr_* is under dual control of the *PhoBR* operon as well as the *RstBA* operon. Both PhoB and RstA binding are known to affect acid sensitivity (Baek & Lee, 2006; Ogasawara et al., 2007; Deliene, 1999).

**Figure 1.**
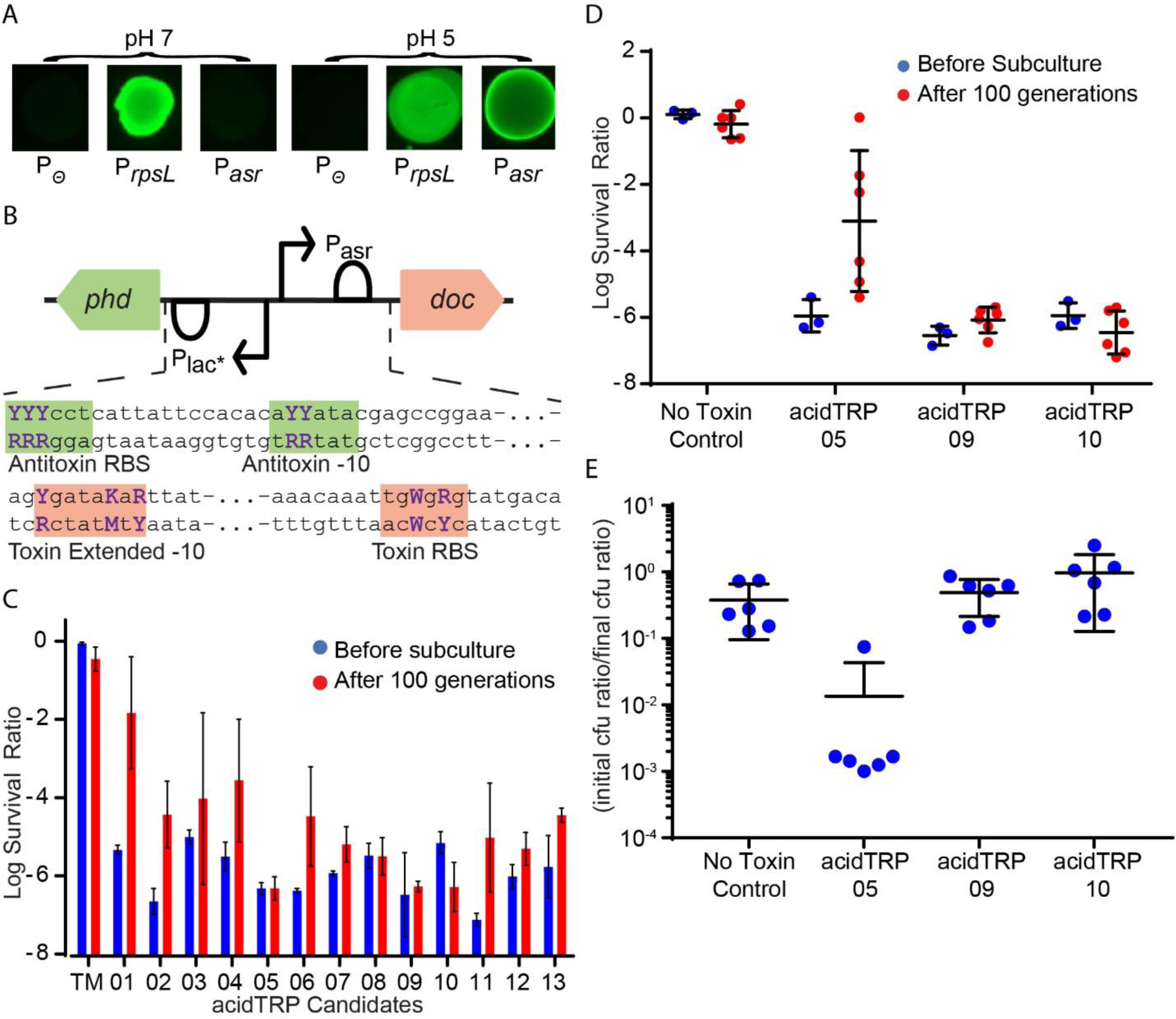
**Construction and screening of a pH sensitive kill switch library** 1A) Expression of native P*_asr_*-GFP at pH 7 vs pH 5. Left to right in each set: no promoter:GFP (negative control), P*rpsl*:GFP (positive control), P*_asr_*:GFP. Images are from the same picture, with comparable levels of bacterial growth. 1B) Structure of acidTRP genetic circuit and degenerate base locations. Black arrows: promoter elements, semi circle: ribosome binding site, coloured arrows: genes. The sequence of the antitoxin and toxin regulatory regions are shown with the RNAP −10 binding sites and ribosome binding sites highlighted (antitoxin green, toxin red). Purple bases indicate degenerate locations (Y = C/T, K = G/T, R = A/G, W = A/T). P*lac\** indicates a constitutive lac promoter. 1C) Survival ratio for acidTRP candidates in *E. coli* TB10 before (blue lines) and after (red lines) 100 generations of growth. TM = toxin mutant control. Error bars represent the standard deviation from two biological replicates. 1D) Survival ratio for candidates AT05, AT09 and AT10 in *E. coli* K-12 MG1655 before (blue) and after (red) 100 generations of growth. 3 technical replicates where used for the before data points and 6 biological replicates were used for the after data points. Error bars represent the standard deviation. 1E) Competitive growth assay between acidTRP strains and *E. coli* K12 MG1655. Mixed cultures with approximately equal titers of an acidTRP strain and MG1655 were grown at pH 7 for around 70 generations. (Initial ratio of acidTRP candidate to MG1655)/(final ratio of acidTRP candidate to MG1655) is plotted for co-cultures of ATTM, AT05, AT09 and AT10 each with MG1655. For candidate 05, the bottom 5 data points represent the limit of detection as no acidTRP-05 colonies were observed after co-culture with MG1655. Data points are 6 biological replicates. Error bars are standard deviation.

In order to maintain stable control over a bacterial population, a containment system should not lose its function due to mutations over multiple generations. To achieve this, a toxin/antitoxin system was used, in which the antitoxin is constitutively expressed at a low level to mitigate any detrimental effect due to leaky expression of the toxin in permissive conditions. We used the type II toxin-antitoxin system Doc/Phd. Doc phosphorylates the elongation factor EF-Tu, inhibiting translation and preventing growth (Castro-Roa et al., 2013; Liu et al., 2008). The antitoxin Phd binds to Doc, inhibiting the kinase activity (McKinley & Magnuson, 2005).

To screen for strains with low escape rates and optimal evolutionary stability, a kill switch library with varying rates of toxin and antitoxin expression was created via rational design. Phd is expressed under a modified constitutive LacUV5 promoter (Malan & McClure, 1984), with Doc expression controlled by P*_asr_*. In a similar technique to the creation of the temperature sensitive kill switch cryodeath (Stirling et al., 2017), degenerate bases were introduced at three locations in the antitoxin RBS and two locations in the antitoxin −10 RNA polymerase binding site, as well as at two locations in the toxin RBS and three locations in the toxin −10 RNA polymerase binding site (Figure 1B). Each of these ten modifications can result in one of two possible nucleotides, giving a library size of 2^10^ = 1024 possible combinations. A construct was made with a frameshift mutation in the toxin ORF to act as a negative control and named acidTRP toxin mutant (ATTM).

The Kill Switch Library was screened to identify candidates that had a low survival ratio and did not lose their function across generations. The library was transformed into the *E. coli* strain TB10 (Caulcott & Rhodes, 1986; Johnson et al., 2004) using the lambda red recombineering genes (Poteete & Fenton, 2000). 200 Individual colonies were re-streaked on pH 7 and pH 5 plates, 41 of which displayed a notable growth defect at pH 5. These colonies were then grown in pH 7 liquid media and plated on M9 buffered to either pH 7 or pH 5. The CFU at each pH was compared to determine a survival ratio for these candidates, and those with a survival ratio of less than 1 in 10^5^ were identified as potential candidates (Figure 1C, blue bars). These candidates were named acidTRP-01 through 13 (AT01-AT13). Next, to assess evolutionary stability, each candidate was passaged at pH 7 for over 100 generations. Candidates AT05, AT09 and AT10 all showed survival ratios of 10^-6^ or less after this growth period (Figure 1C, red bars) and were selected for further analysis. Each was transferred into the *E. coli* K12 strain MG1655 via P1 transduction, and the evolution experiment was repeated with more biological replicates (Figure 1D). Whilst candidates AT09 and AT10 showed no discernable difference before and after a period of growth, AT05 had a noticeable increase in survival ratio.

In permissive conditions, candidates AT09 and AT10 showed no disadvantage in growth rate when compared to MG1655. Candidates AT05, AT09 and AT10 were each co-cultured with MG1655 in pH 7 media and passaged for approximately 60 generations, the population ratio of each kill switch strain to MG1655 was measured by comparing the CFUs before and after growth across 6 biological repeats (Figure 1E). Candidates AT09 and AT10 showed no significant difference in the ratios of kill switch strain to MG1655 before and after the period of growth when compared to a co-culture of the toxin mutant control and MG1655. For candidate AT05, the kill switch strain was consistently out-competed by MG1655.

With no discernable difference between the survival ratio or evolutionary stability of AT09 and AT10, AT09 was chosen for further experiments. To characterise an induction curve, GFP was expressed under promoter Pasr-AT09. Cultures were grown to mid log phase at a range of pHs, and GFP fluorescence was measured using flow cytometry. Induction began at pH 6, with over 6000-fold increase in GFP production between pH 6 and pH 4.4 (Figure S1).

AT09 was combined with cryodeath (Stirling et al., 2017) resulting in a multiplicative effect on survival ratio in non-permissive conditions. Cryodeath is a temperature sensitive kill switch that uses the regulatory system from cold shock protein A (*cspA*) and the toxin antitoxin system CcdB/CcdA. At 37 °C the toxin is repressed, but at 22 °C an increase in expression of the toxin leads to population death with a survival ratio of around 10^-5^. The combined strain, containing both cryodeath and acidTRP (CD-AT) was grown at 37 °C in pH 7 media and then plated on both pH 7 and pH 5 plates, which were incubated at either 37 °C or 22 °C. Strains with a frame-shift mutation in one or both of the toxins were assayed in parallel as controls. Whilst the strain with individual kill switches behaved as expected, with survival ratios of 10^-5^ (only cryodeath functional) to 10^-6^ (just acidTRP functional), the double kill switch strain displayed a survival ratio that could not be detected as no colonies were formed when plating on pH 5 media and incubating at 22 °C (Figure 2A), even when 10^11^ cfu was plated.

**Figure 2.**
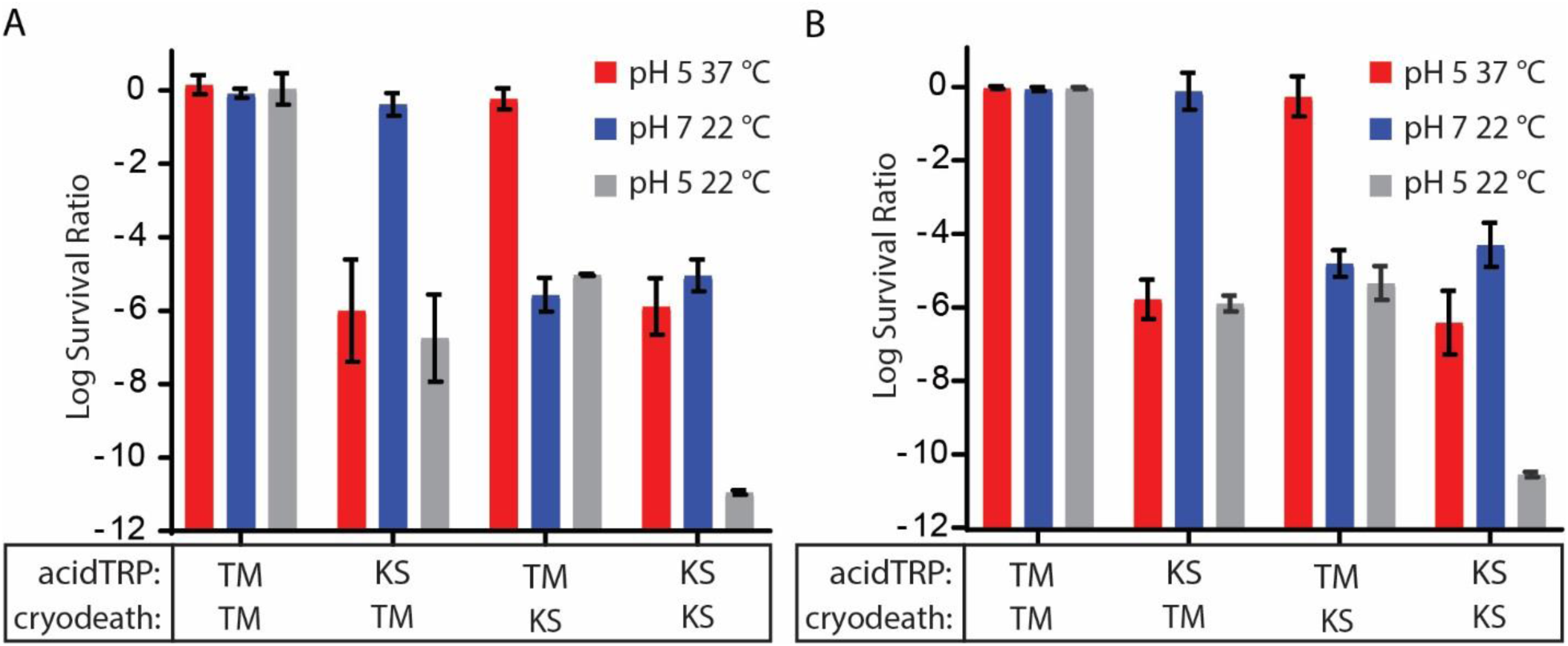
**Combining the pH sensitive kill switch acidTRP and the temperature sensitive kill switch cryodeath (Stirling et al., 2017) into a single strain.** 2A) The combination of acidTRP and cryodeath kill switches gives a multiplicative reduction in survival. Four strains with either a kill switch (KS) or toxin mutant (TM) inserted into the acidTRP and cryodeath loci were cultured in pH 7 media at 37 °C to late log phase. A dilution series was then plated on both pH 7 and pH 5 plates and grown at both 37 °C and 22 °C. Survival ratio for each strain was calculated when comparing the cfu at 37 °C/pH 5, 22 °C/ pH 7 and 22 °C/ pH 5 to the cfu at 37 °C/ pH 7. No colonies were observed on 22 °C/pH 5 plates for the double kill switch strain, so data points represent the limit of detection (ie total cfu on 37 °C/ pH 7 plate). Error bars represent the standard deviation from two technical replicates. 2B) The combined kill switch strain is evolutionarily stable. The experiment from Figure 2A was repeated after a growth period of approximately 100 generations. Error bars represent the standard deviation from 6 biological repeats.

CD-AT did not lose its capacity for containment after 100 generations of growth. CD-AT was passaged at 37 degrees for approximately 100 generations, along with the toxin mutant controls. Survival ratios for all strains were comparable to before the period of evolution (Figure 2B).

### A novel counter circuit, responding to the falling edge of a stimulus pulse

We next constructed a counter circuit to allow our sensors to record multiple discrete pulses of a given stimulus. We built upon our previous design of a bacterial memory circuit, which is based on the OR operon of bacteriophage λ, and which is capable of sensing and recording transient stimuli (Kotula et al., 2014; Naydich et al., 2019; Riglar et al., 2017). We created a two-counter circuit (Figure 3A), based on the OR operon of bacteriophage 434, which produces a fluorescent response only after sensing two distinct pulses of a stimulus (e.g., low pH). Importantly, this circuit’s ability to count pulses is not sensitive to an extended duration or concentration of the applied pulse. This feature is achieved by delaying the first count until the first stimulus pulse is removed (i.e., until the falling edge of the pulse).

**Figure 3:**
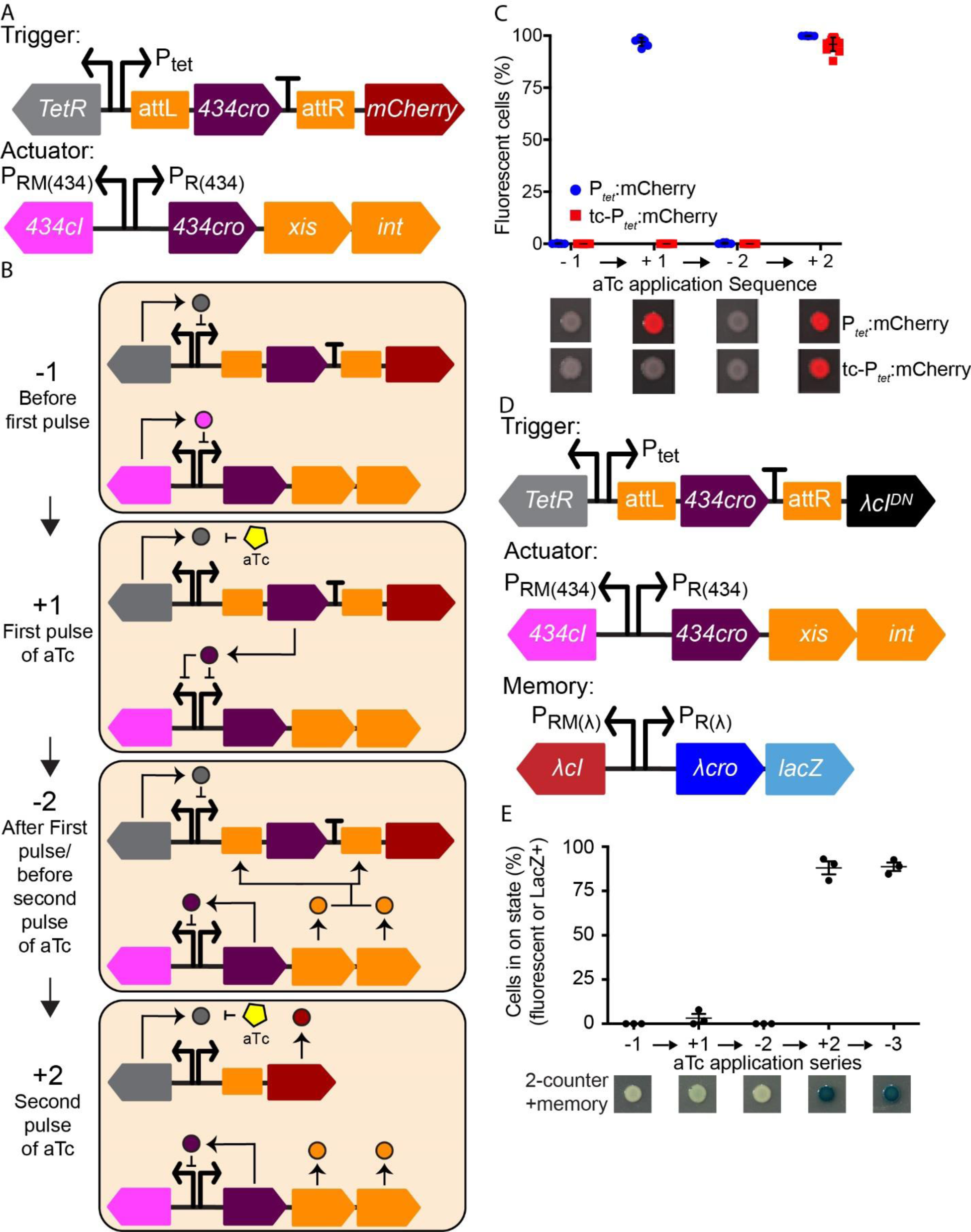
genetic circuits for 2 different two-counter circuits using an actuator based on phage 434 cI and Cro transcription factors. Black arrows: promoter elements, attL/attR: 434 Xis/Int attachment sites, T: terminator, coloured arrows: genes. 3A) Schematic of the 434 two-counter, including trigger and actuator elements. 3B) Schematic of trigger response and modification during two subsequent pulses of aTc. −1: actuator in cI state, TetR represses expression on the trigger. +1: aTc represses TetR, Cro is expressed from the trigger and represses both cI and Cro expression from the actuator. −2: Actuator in Cro state. expression of Xis and Int targets attachment sites on the trigger, excising *cro* and the terminator. +2: aTc represses TetR, mCherry is expressed from the trigger. Continued expression of Xis and Int prevents reintegration of excised DNA. 3C) Response of the 434 two-counter circuit (n = 7) to an aTc induction time course with two stimulus pulses (red). aTc (100 ng/ml) was applied to liquid culture and washed out for 4 hour growth periods. A P*_tet_*-mCherry strain (n = 4) is tested as a control (blue). Error bars represent SD. Spots of cultures on agar media (with or without aTc) are shown below at each step of the time course. Spot images consist of an mCherry color overlay on a grayscale brightfield image. 3D) A 3-element Two-counter circuit that activates a memory element to register and record two exposures to aTc. 3E) Response of 434 two-counter memory circuit and 434 two-counter circuit to an aTc induction time course with two stimulus pulses, followed by a final growth step in the absence of stimulus. aTc (100 ng/ml) was subsequently applied and washed out for growth periods of at least 4 hours. Error bars represent SD of three biological replicates. Spots of cultures on agar media containing X-gal (with or without aTc) are shown below at each step of the time course, full-color brightfield image.

The two-counter circuit consists of two constructs: a trigger, which senses a stimulus, and an actuator, which modifies the trigger after the first stimulus pulse is removed, thus enabling a different response to the second pulse. In our initial circuit, the trigger was designed to respond to anhydrotetracycline (aTc) using a P*_tet_* promoter. The actuator is based on a bi-stable switch derived from the phage 434 OR operon, and its two states correspond to the mutually-repressive phage 434 proteins, Cro (under control of PR) and cI (under control of PRM) (Figure 3A). In our two-counter, the actuator begins in the cI state, with the PRM promoter active. In this state, cI represses PR, which controls the production of Cro and the λ Xis and Int proteins (Figure 3B, −1 step). When aTc is applied, Cro is produced from the trigger, while the terminator after the *cro* gene ensures that mCherry is not transcribed. The high level of Cro produced from the trigger in the presence of aTc places the actuator in a sustained intermediate state, with both PRM and PR repressed by Cro, due to the ability of Cro to repress PR at high concentrations (Figure 3B, +1 step). When aTc is removed (the falling edge of the pulse), Cro is no longer produced from the trigger, and the level of Cro in the cell falls, leading to derepression of PR. This, in turn, leads to expression of Cro from the PR promoter, which maintains repression of PRM. It also leads to the production of Xis and Int, which excise the sequence between attL and attR in the trigger, leaving an attB scar site (Figure 3B, −2 step). As a result of this excision, the trigger is primed for the second count, and a second aTc induction leads to the expression of the mCherry reporter. The continued presence of Xis prevents further re-integration (Abremski & Gottesman, 1982; Better et al., 1983) (Figure 3B, +2 step. −1 = before first pulse of aTc, +1 = during the first pulse of aTc, −2 = After first pulse/before the second pulse of aTc, +2 = during the second pulse of aTc.

The two-counter strain was tested using the P*_tet_* promoter controlling the trigger and applying a time course of sequential pulses of aTc induction and quantifying the response of the strain via mCherry fluorescence (Figure 3C). The two-counter showed robust counting behavior, with two pulses required to produce mCherry fluorescence. There was virtually no mCherry signal during or after the first aTc pulse, even when the pulse was applied for up to 24 hours. At the second aTc application, a high percentage of cells displayed fluorescence (-aTc1, +aTc1, and −aTc2: 0.0% ± 0.0% SD; +aTc2: 95.9% ± 3.3% SD; n = 12). The excision of the two-counter trigger was confirmed by PCR amplification of the trigger region at each step of the time course. Gel electrophoresis of the PCR product showed a reduction in trigger length consistent with excision of the region between attL and attR (full trigger length: 4.0 kb; excised trigger length: 3.3 kb) (Figure S3A). Furthermore, analysis by PCR confirmed that any mCherry-negative colonies had not undergone excision and still had the original trigger.

The two-counter also displays a high signal-to-noise ratio (Figure S3B) due to the stability of the phage 434 lysis–lysogeny switch in the cI state. The 434 cI protein used to maintain the memory-off state contains an ind-mutation, based on ind-mutants in phage λ (Gimble & Sauer, 1985). This mutation prevents RecA-mediated cleavage—the typical mechanism of induction in wild-type lambdoid phages during an SOS response (Sauer et al., 1982) —allowing it to stably maintain the cI state prior to the first application of aTc.

The bistable nature of the cI–Cro switch also produces a digital output with a clear demarcation in fluorescence between mCherry-negative and mCherry-positive cells (Figure S3B). Prior to the second application of aTc, all cells maintained a fluorescence below ∼100 a.u. At the second application of aTc, a bimodal population was observed, in which there was a marked increase in mCherry fluorescence for cells that have completed trigger excision as a result of the removal of the first aTc stimulus. Because the population of cells that have experienced two pulses does not overlap with the population of cells experiencing fewer than two pulses, the readout of the count can be inferred at a single-cell level. In addition to the aTc-responsive two-counter based on the phage 434 OR operon, we also constructed a similar circuit based on the OR operon from phage lambda (Figures S3C-F). These results demonstrate our ability to construct falling-edge pulse counters based on lambdoid phage switches with high stability and signal-to-noise ratio.

A key motivation for constructing sensing circuits in bacteria is the potential to deploy them in inaccessible environments, such as the mammalian gut. For instance, bacteria containing a pulse counter might be used to track the number of times they encounter a signal during transit through the intestine. After exiting the gut, the bacteria may be recovered to obtain a non-invasive readout of the registered count. While the two-counter described above produces distinct responses to two discrete pulses of a stimulus, its reporter readout requires active presence of that stimulus, as is the case with previous pulse counter circuits (Friedland et al., 2009). For a deployment-and-recovery scenario, a circuit must be able to record the count and produce a sustained readout which is interpretable even when the sensor is removed from the sensing environment.

We constructed a two-counter memory circuit (tc-P*_tet_*-memory), which allows both sensing and recording of two discrete stimulus pulses (Figure 3D). The tc-P*_tet_*-memory consists of a trigger, an actuator based on the phage 434 cI–Cro switch, and a memory switch based on the phage λ cI–Cro switch (Kotula et al., 2014; Naydich et al., 2019; Riglar et al., 2017). With the first aTc application, the circuit behaves identically to the 434 two-counter, producing Xis and Int from the actuator with the falling edge of the first pulse. On the second aTc application, the trigger produces λ cIDN (Naydich et al., 2019), which flips the λ memory switch to the Cro (memory-on) state so that it continuously produces a LacZ reporter. As with the previous two-counter, the two-counter memory circuit is designed to respond to the falling edge of the first stimulus pulse and the rising edge of the second.

The two-counter memory strain was subjected to a time course of aTc pulses, with a third aTc-free step after the second aTc pulse. The response at each step was quantified by plating cultures on X-gal indicator plates (+/−aTc, as appropriate) (Figure 3E). The strain showed nearly zero activation prior to the second pulse of aTc. The second aTc application resulted in a high percentage of cells in the on state, which persisted even after aTc was removed (-aTc1: 0.0% ± 0.0% SD; +aTc1: 3.2% ± 4.1% SD; -aTc2: 0.0% ± 0.0% SD; +aTc2: 88.0% ± 6.3% SD; -aTc3: 88.7% ± 4.1% SD; n = 3). The two-counter memory circuit demonstrates our ability to incorporate complex counting and recording logic by combining multiple lambdoid phage switches in a synthetic context.

### pH sensitive counting and containment mechanisms

The pulse counting systems developed for P*_tet_* were reconstructed with P*_asr_* controlling expression of *cro* and *gfp* (Figure 4A), yielding similar results. A plasmid with the P*_asr_* promoter from acidTRP-09 controlling expression of a trigger element with GFP as the final reporter gene was transformed into an *E. coli* K12 strain containing the actuator, creating strain tc-P*_asr_*-AT09:GFP (Figure 4A). This strain was assayed by applying a time course of 5 hour growth steps alternating between pH 7 and pH 5. There was virtually no GFP signal before, during or after the first exposure to pH 5. Upon the second exposure to pH 5, almost 100% of cells displayed fluorescence (-pH5 1: 0.0% ± 0.0% SD, +pH5 1: 0.05% ± 0.03% SD, -pH5 2: 0.18% ± 0.04 SD; +pH5 2: 98.83 % ± 0.82% SD; n = 3) (Figure 4B).

**Figure 4.**
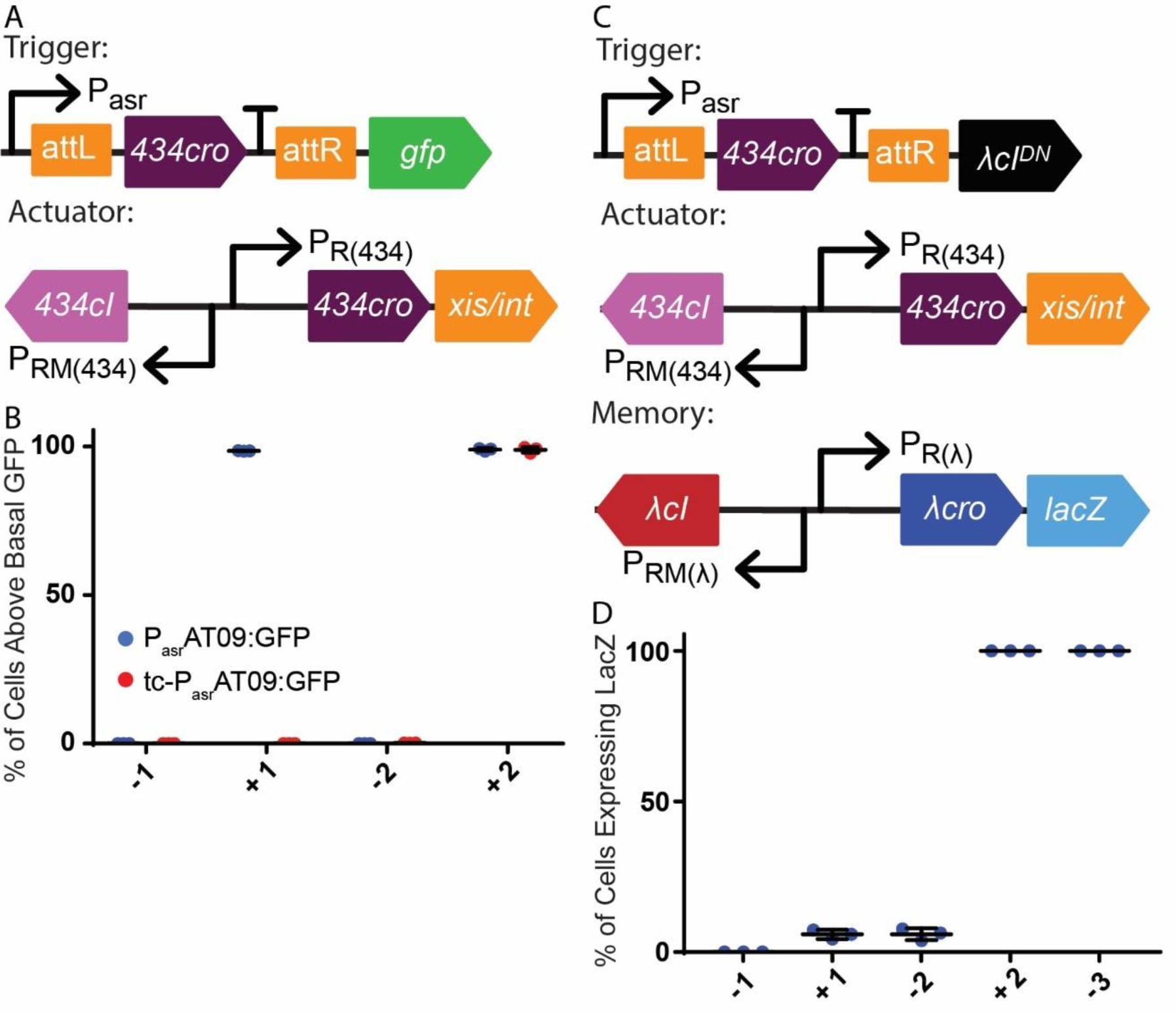
Analysis of the pH sensitive two counter (tc-P*_asr_*AT09:*gfp*) and two counter memory (tc-*Pasr*AT09-memory). Black arrows: promoter elements, attL/attR: 434 Xis/Int attachment sites, T: terminator, coloured arrows: genes. 4A) Schematic of trigger and actuator of tc-P*_asr_*AT09:*gfp* strain. 4B) percentage of cells expressing GFP above basal levels for P*_asr_*AT09:*gfp* (blue) and tc-P*_asr_*AT09:*gfp* (red). −1 = before first exposure to pH 5, +1 = during first exposure, −2 = after first exposure/ before second exposure, +2 = during second exposure. Data points from three biological replicates. Error bars represent the standard deviation. 4C) Schematic of the trigger, actuator and memory cassette of tc-P*_asr_*AT09-memory strain. 4D) The percentage of cells expressing lacZ for tc-P*_asr_*AT09-memory strain. −1 = before first exposure to pH 5, +1 = during first exposure, −2 = after first exposure/before second exposure, +2 = during second exposure, −3 = after the third exposure.

Sustained exposure to pH 5 and pH 7 did not cause unwanted expression of GFP. tc-P*_asr_*AT09:GFP was cultured overnight in pH 7 media, and then exposed to three five-hour growth steps in every possible permutation of pH 7 and pH 5 media (Figure S4A). Even when cultured in pH 5 media for 15 hours (approximately 30 generations), negligible GFP expression was observed. GFP expression was limited to only the culture that had been exposed to pH 5 conditions for two nonconsecutive growth steps (Figure S4B).

A pH sensitive two-count memory circuit using LacZ as a reporter was constructed (tc-P*_asr_*AT09-memory) (Figure 4C). This strain was subjected to a time course with varying pH levels, after which memory response was quantified by plating on pH 7 X-gal indicator plates. Approximately 5% of cells showed switching before the second exposure to pH 5, compared to effectively 100% after the second exposure (Figure 4D).

The pH-sensitive kill switch was combined with the pulse counter system to produce a kill switch that would only produce toxin upon a second exposure to low pH (tc-acidTRP-09). The toxin, *doc*, was inserted in place of a reporter on the trigger element, and the antitoxin, *phd*, was placed upstream of P*_asr_* (Figure 5A). After a single exposure to pH 5 and subsequent growth at pH 7, *434 cro* is excised, and the trigger element of the resulting circuit becomes site between P*_asr_* and the toxin RBS (Figure 5B).

**Figure 5.**
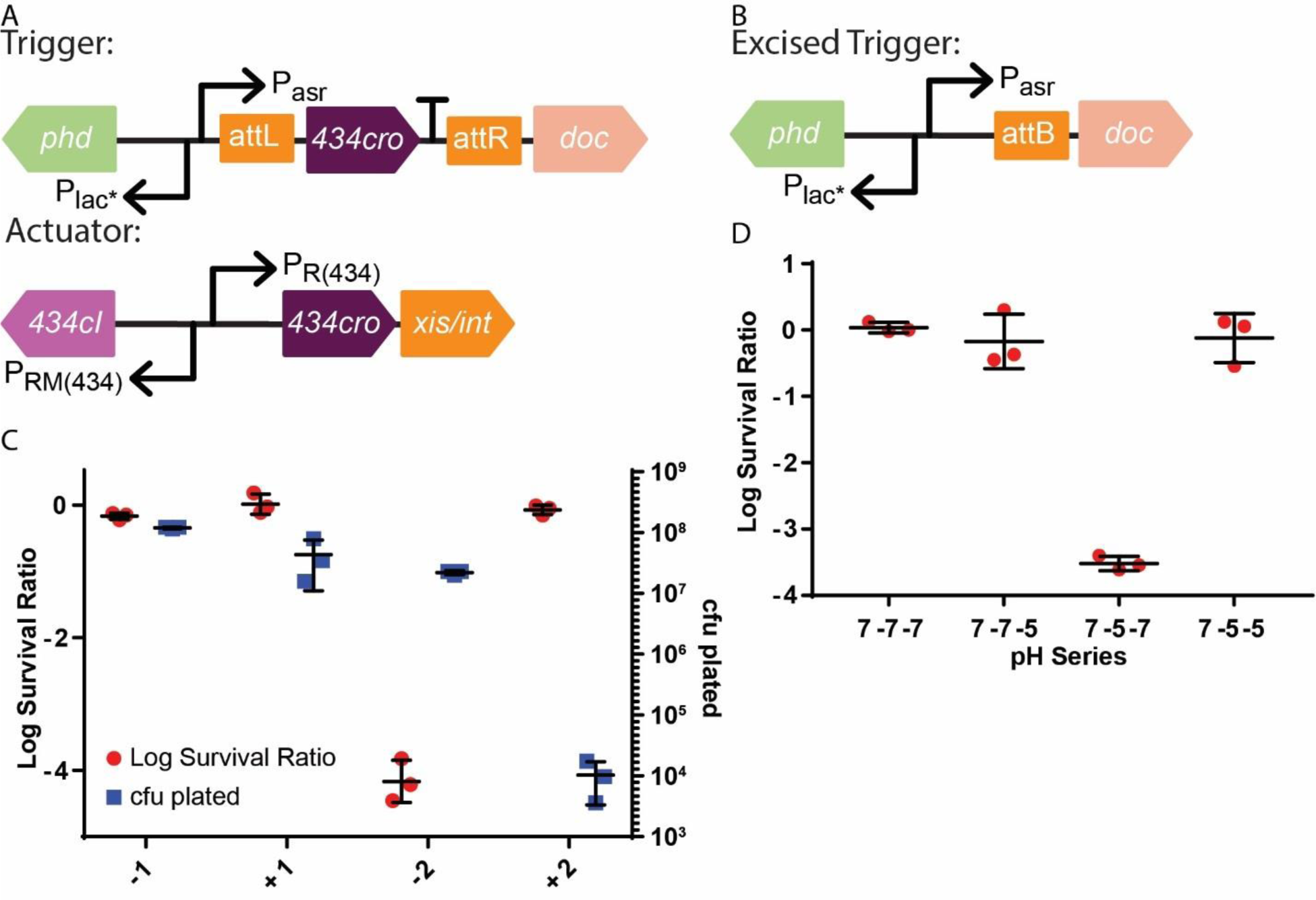
analysis of the two count pH sensitive kill switch (tc-acidTRP-09). Black arrows: promoter elements, attL/attR: 434 Xis/Int attachment sites, T: terminator, coloured arrows: genes. 5A) Schematic of the trigger and actuator of tc-acidTRP-09. 5B) Schematic of the trigger of tc-acidTRP-09 after excision of 434cro element. 5C) Survival ratio (red) and total cfu on pH 7 plates (blue) of tc-acidTRP-09 throughout a time course experiment of varying pH conditions. −1 = before first exposure to pH 5, +1 = during first exposure, −2 = after first exposure and +2 = during second exposure. 5D) Log survival ratio of tc-acidTRP-09 after exposure to 5 hours at 37 °C in three consecutive conditions. pH 7 followed by either pH 7 or pH 5 followed by either pH 7 or pH 5. After exposure to the final condition, cultures were plated on pH 7 and pH 5 plates and a survival ratio was calculated. Only the 7-5-7 culture plated on pH 5 had two, nonconsecutive exposures to pH 5 conditions.

tc-acidTRP-09 displayed a survival ratio of 10^-3^ to 10^-4^ after undergoing a time course of pH 7 and pH 5 conditions. Length of exposure to the different pH levels was explored to identify the induction conditions resulting in the lowest survival ratio, and a pattern of 5 hours at pH 7 (−1), 10 hours at pH 5 (+1), 5 hours at pH 7 (−2) and finally 5 hours at pH 5 (+2) was settled on, all at 37 °C (data not shown for alternate conditions). The initial growth at pH 7 could also be an overnight growth at room temperature without affecting the resulting survival ratio (data not shown). After each growth step, serial dilutions of each culture were plated on both pH 7 and pH 5 plates, and a survival ratio was calculated. After the −1 and +1 steps, the log survival ratio was approximately 0, indicating excision had not taken place and *doc* was not being expressed. After the −2 step, a log survival ratio of approximately −4 was observed, indicating excision had occurred. After the +2 step, any surviving cells were plated, and the log survival ratio had returned to approximately 0. This result is expected, presumably because any cells remaining at the end of the +2 step had survived due to a mutation preventing tc-acidTRP-09 from functioning or because they had not performed trigger excision during the previous step—or a mixture of both (Figure 5C). Accordingly, the total cfu in the culture at the end of the +2 step was dramatically reduced (Figure 5C), indicating only a small proportion of the population was still viable. Surprisingly, PCR evidence showed that 100% (n=7) of escapees on pH 5 plates had excised, suggesting that toxin mutation was the primary reason for escape in this context.

tc-acidTRP-09 only displayed a decrease in survival ratio when exposed to pH 5 conditions on two separate, non-consecutive occasions. After an initial growth at pH 7, tc-acidTRP-09 was grown for at least 5 hours at either pH 7 or pH 5 for two consecutive growth steps in each possible permutation, represented by the 2^nd^ and 3^rd^ growth steps in Figure S4A. A serial dilution of the resultant cultures was then plated on pH 7 and pH 5 plates, and a survival ratio was calculated for each set of conditions (Figure 5D). Sustained exposure to pH 5 and pH 7 failed to drop the survival ratio below 0, whereas a single exposure to pH 5 followed by a growth period at pH 7 dropped the survival ratio to approximately 10^-3.5^. This is expected, as growth on the pH 5 plate would be the second exposure to pH 5 conditions and would therefore induce expression of the toxin only in strains that had excised the *cro* element.

## Discussion

We have used the *E. coli* pH sensitive promoter P*_asr_* to construct toxin/anti-toxin based containment system called acidTRP. This containment system, which uses a kill switch activated at low-pH conditions, was shown to be evolutionarily stable for at least 100 generations and to result in a survival ratio of less than 10^-6^ upon exposure to pH 5. To achieve these attributes, we constructed and screened a rationally designed toxin/antitoxin library with varying levels of expression dictated by degenerate bases in the promoter RNAP −10 and ribosome binding sites of both the toxin and the antitoxin. This method was previously used to build the temperature sensitive kill switch cryodeath (Stirling et al., 2017), and the successful construction of acidTRP demonstrates the broader application of the method.

AcidTRP was successfully combined with cryodeath into a single strain that responded to exposure to both low pH and low temperature. Respectively, each containment system conferred a survival ratio of 10^-6^ and 10^-5^. The combined strain displayed a survival ratio below the limit of detection of 10^-11^, indicating the two systems combined in a multiplicative manner. This is to be expected, as cryodeath uses an orthogonal type II toxin/antitoxin system, CcdB/CcdA, with a different cellular target (Madl et al., 2006; Wright et al., 2013), limiting interaction between the two kill switches and insuring that an individual cell must contain disabling mutations for both systems in order to survive. In addition, it has been shown that there is little cross talk between the regulation of P*_asr_* and the cold shock response promoter P*cspA* used in cryodeath (Hoynes-O’Connor et al., 2017).

For effective control of a bacterial population, a containment system’s survival ratio must be low enough that the probability of an escape event over the time period of the bacterial population’s intended use is negligible. In the example of the human digestive system, an individual expels *E. coli* at a rate of approximately 10^7^ cfu per gram of excrement (Farnleitner et al., 2011) and produces approximately 300 g of excrement per day (Hosseini, 2000). Therefore, for a population of *E. coli* residing in the human digestive tract over the course of one year, a survival ratio of less than 10^-11^ would result in fewer than 10 escape events. Whilst this does not take into account any potential fitness defect for the bacteria accrued in the process, the current standard for containment of transgenic *E. coli* host vectors is set by the NIH as no more than 1 in 10^8^ surviving to pass on genetic code under specified conditions (NIH Guidelines, 2019, Apendix I-I-B). Whilst our containment system meets this standard, it is possible a more stringent standard is required for applications of *E. coli* outside of a laboratory.

In addition to the combined pH and temperature sensitive kill switches, we constructed pulse-sensing and pulse-counting circuits, based on a bi-stable, bacteriophage-derived switch with a built-in negative feedback mechanism. Uniquely for synthetic biological circuits, we have shown that our counters can respond to the falling edge of a stimulus pulse, a critical feature for circuits designed to produce unique responses to multiple discrete pulses of the same stimulus.

Our circuits demonstrated robust counting behavior *in vitro*, exhibiting three distinct advantages over previous counters. First, our counters record only a single count for a single pulse, even when pulses were applied for up to 15 hours across multiple back dilutions (Figures S4B and 5D). Because the sustained intermediate state of the cI–Cro switch can be maintained indefinitely while the trigger is activated, we posit that there is no upper limit to the concentration and duration of an applied pulse that would cause the circuit to record more than a single count. Second, our counters have a tightly controlled off-state, ensuring no spontaneous switching, resulting in a high signal-to-noise ratio (Figures 3C, 3E, 4B, 4D, 5C and 5D). Third, the counters are digital reporters, which enables an interpretable readout of the count on a single-cell basis. This improves upon previous pulse counters, which produced reporting states that overlap between populations experiencing varying numbers of pulses, and thus were only interpretable when sampled as a population (Friedland et al., 2009). Single-cell readable counts are particularly important for applications in which there may be a limited number of counter cells recovered from the environment, and therefore no ability to analyze population-level signals.

By combining two phage-derived bidirectional switches in a single circuit, we created a two-counter with memory, which continuously produces a reporter after a cell has experienced the falling edge of a first pulse followed by the rising edge of a second. Like the 434 and λ two-counters, this circuit showed efficient counting *in vitro* (Figures 3E and 4D).

The ability to reliably respond to distinct stimulus pulses enables a range of potential applications in tracking and responding to extracellular and intracellular signals. We have shown the counter can easily be adjusted to respond to a variety of sensors by altering the promoter controlling expression of *cro* on the trigger, such as by using the P*_asr_* promoter. Synthetic biology has taken advantage of natural and designed parts to construct complex circuits, which incorporate increasingly sophisticated logic. Our demonstration of falling-edge pulse counters expands the repertoire of synthetic circuit functions and provides a scalable design for higher-order sensing, while our integration of kill switch and counter circuits demonstrates the use of such sensing to program specific cellular behaviors.

## Acknowledgements

We owe a lot to the dedicated work of Loden Dundutsang, producing the many hundreds of pH buffered plates this work required. We thank David Riglar for consultations on designing and implementing the pulse counting systems. This work was supported by Defense Advanced Research Projects Agency grant [HR0011-15-C-0094] and funds from the Wyss Institute for Biologically Inspired Engineering. F.S. acknowledges funding from NIH training grant [5T32GM007598]. A.N. acknowledges funding from National Science Foundation Graduate Research Fellowship [DGE1144152 and DGE1745303].

## Author Contributions

Conceptualization, F.S., A.N., M.C, H.W. and J.W.; Investigation, F.S., A.N., J.B., R.B., M.C., H.W., E.R., S.O., S.G., A.C., Writing – Original Draft, F.S., A.N., Writing – Review & Editing, F.S., A.N., J.W., P.A.S., Funding Acquisition, P.A.S. and J.W.

## Methods section

### EXPERIMENTAL MODEL AND SUBJECT DETAILS

#### *E. coli* K12 strain MG1655

Used as the basis for all strains except for the tc-memory strains (both aTc and pH sensitive). Maintained using established protocols for *E. coli*. Strains that contained acidTRP were maintained in buffered pH 7 media, unless being assayed, and were prevented from growing above OD 1.5 to prevent acidification of the media. Strains that contained cryodeath candidates were maintained at a constant temperature of 37 °C unless being assayed.

#### *E. coli* K12 strain TB10

A derivative of MG1655, with a large section of the lambda prophage genome inserted into the biotin operon. Used to recombine the acidTRP library and in strain construction for two counter circuits. cI has been mutated to the temperature sensitive variant, cI^857^, allowing for temperature sensitive induction of the lambda red genes. Maintained using established protocols for E. coli, except all growth was kept at 30 °C.

#### *E. coli* strain NGF1

*E. coli* NGF-1 is a strain originally isolated from the murine gut, which has proven to be an efficient and persistent colonizer and a reliable platform for the deployment of engineered circuits (Kotula et al., 2014).

### METHOD DETAILS

#### Media and growth conditions

Unless otherwise specified LB media with relevant antibiotic was used for all growth conditions. For the Buffered media, M9 minimal media supplemented with 1 mM MgSO4, 0.4% w/v glucose, 100 mM CaCl2 and 1 gram per liter of yeast extract (Difco) was used. For the pH 7 media the 5X M9 salts consisted of 15 mg/L CaCl2, 5 g/L NH4CL, 2.5 g/L NaCl, 30 g/L Na2HPO4, 15 g/L KH2PO4. For the pH 5 media the 5 x M9 salts consisted of 15 mg/L CaCl2, 5 g/L NH4CL, 2.5 g/L NaCl, 284 mg/L Na2HPO4, 43.4 g/L KH2PO4. Both were titrated to the desired pH using HCl or KOH. It should be noted that the pH 7 buffered media had a noticeably better buffering capacity. For the induction curve HCl or KOH where added to the pH 5 media (final pH 4.4–5.8) and the pH 7 media (final pH 6). To make plates, 15 g/L of bacto agar was added. For all buffered media, distilled water was used, and it was observed that MillliQ ultrapure water resulted in minimal growth, presumably due to the lack of trace elements in the resultant media. Unless otherwise stated, in all instances of antibiotic use the following concentrations were employed: Amp 100 μg/ml, Kan 50 μg/ml, Gent 10 μg/ml, Strep 100ug/ml, Cam 25 μg/ml. Unless otherwise specified, P*_tet_* trigger strains were induced by adding 100 ng/ml aTc to liquid media or to agar plates. To quantify two-counter memory response on indicator plates, agar was supplemented with X-gal (60 µg/ml).

#### Acid sensitive reporter plasmid design

All primer sequences are given in Table S5. The P*_asr_* regulatory region was ordered as reagents for a BioXP from SGI DNA, along with homology to the plasmid pUA66 (Table S1). pUA66 was linearised with the primers TS1 and TS2 and combined with the P*_asr_* fragment using an NEB Gibson assembly kit according to the manufacturers guidelines, to create a plasmid with pH sensitive expression of GFP. The resultant plasmid was transformed into electrocompetent MG1655 cells.

#### Construction of the acidTRP library

The acidTRP library was ordered from SGI DNA as a custom order, with the 10 degenerate bases included (Table S2). On each end are 100 bp of homology to TrpC for homologous recombination into the genome. The resultant product was amplified using primers LS17 and LS18. After 35 cycles, the reaction was passed through a zymo clean and concentrator kit, and then run for 5 additional cycle with new PCR reagents. This step is to ensure that the final product contained two strands that matched completely, as mismatched strands may undergo mismatch repair in a strand independent manner, resulting in double stranded breaks as gap-repair polymerases meet.

#### P1 transduction

P1 lysates were constructed according to previously published methods (L. C. Thomason et al., 2007). Overnight cultures of donor strains were grown at relevant temperatures in LB, before being back diluted into 50 fold into 5 mL of LB supplemented with 0.2% w/v glucose, 20 mM MgCl2 and 5 mM CaCl2. After approximately 45 minutes of growth at the relevant temperature, between 10 and 100 ul of high titer (pfu > 100) lysate was added and the culture was grown until lysis had occurred. Lysates were then centrifuged at 4000 rpm for 5 minutes and the supernatant filtered using a 0.2 micron filter. Lysates were stored for up to 6 months at 4C.

#### Recombineering

Electrocompetent cells were prepared for transformation using previously published methods (L. Thomason et al., 2007). The acidTRP library was transformed into the recombinant strain TB10 that had been induced for 15 minutes in a 42 °C water bath with manual shaking every two minutes. Approximately 100 ng of DNA was combined with 50 ml of cells in a 0.1 cm cuvette, and electroporated using EC1 setting on a Biorad Micropulser. Cells were recovered in SOC medium for one hour before being spread on pH 7 buffered plates with kanamycin.

#### Initial acidTRP library screen

Approximately 200 colonies from the recombination of the library were struck out on both pH 7 and pH 5 plates. Any colony that showed a growth difference between the two conditions was struck to single colonies on both conditions, using cells from the pH 7 plate. Colonies that still displayed a growth difference between the two conditions were used to inoculate pH 7 liquid media and grown for 4 hours at 37 °C, and then used to make 25% glycerol stocks. All candidates were sequenced to determine the identity of each degenerate base (Table S3).

#### Constructing the toxin mutant control

The primer pairs LS17 and LS26 along with LS25 and LS18 were used to PCR the acidTRP library fragment from SGI DNA. The two resultant fragments were combined using an NEB Gibson assembly kit. The resultant product was transformed into TB10, and a single colony was picked to assay for lack of sensitivity to pH before being sequenced to confirm the frame shift mutation and identify the degenerate bases (Table S3).

#### pH sensitive survival assay

Glycerol stocks of acidTRP were used to inoculate pH 7 media and grown at 37 degrees until late log phase, approximately OD 1.0. One-hundred-fold serial dilutions of each culture were made in pH 7 media and plated onto both pH 7 and pH 5 buffered plates. Plates were then incubated for at least 48 hours at 37 °C. The cfu on each plate was counted and a ratio of pH 5 cfu to pH 7 cfu was calculated and termed the survival ratio.

#### Evolutionary stability of acidTRP strains

Glycerol stocks of acidTRP strains were used to inoculate 2 ml of pH 7 media, and then grown for approximately 24 hours at room temperature. These cultures were then sequentially passaged for another 20 growth steps by back diluting 100 fold into 2 ml of pH 7 media. OD was measured each day and used to calculate the number of generations that had passed. pH was measured after each growth step to ensure the pH of the media remained approximately at pH 7.

#### Competitive Growth Assay

Overnight cultures of MG1655 mixed with acidTRP-TM, 05, 09 and 10 were grown at room temperature to mid log phase. 100ul of the MG1655 culture was combined with each other culture, and then used to inoculate pH 7 media. Glycerol stocks were made of this initial inoculation. Cultures were passaged once every 24 hours diluting 1:1000 into pH 7 media. After 8 days, glycerol stocks were made of the final culture. Glycerol stocks were then defrosted, and a dilution series was plated onto both LB kanamycin and LB plates. The cfu ratio between the two types of plate was then used to determine the ratio of kill switch strains to MG1655.

#### Combined acidTRP and cryodeath survival assays

A P1 transduction was used to transfer each of the acidTRP-09 and acidTRP-TM cassettes into each of the cryodeath and cryodeath-TM strains, creating 4 new strains (acidTRP-TM/cryodeath-TM, acidTRP/cryodeath-TM, acidTRP-TM/cryodeath and acidTRP/cryodeath). For the first three, survival assays were conducted as before. For the acidTRP/cryodeath strain, glycerol stocks were used to inoculate 500 ml of pH 7 media and grown for ∼ 10 hours at 37 °C until ∼ OD 1. Cultures were centrifuged at 4000 rpm at 4 °C for 30 minutes in a 500 ml bucket centrifuge, resuspended in 20 ml of pH 7 media and transferred to 50 ml falcon tubes then spun down at 4000 rpm at 4 °C for 30 minutes. The supernatant was removed, and the pellet was resuspended in an additional 5 ml of pH 7 media (this represents the undiluted culture, 10^0^). The undiluted culture was then diluted 10 fold followed by an additional 5 100 fold dilutions (10^1^, 10^3^, 10^5^, 10^7^, 10^9^, 10^11^ dilutions). 225 *μl of e*ach dilution was then plated on two pH 7 and two pH 5 plates, and a plate of each pH was grown at both 37 °C and room temperature (∼22 °C) for at least 5 days. 225 *μl* of the undiluted, 10^0^ culture was also plated on 10 different pH 5 plates and grown at room temperature for the same time period. cfus were then counted in all conditions and a ratio of each compared to pH 7/37 °C was calculated as the survival assay for each condition.

#### Evolutionary stability of combined acidTRP and cryodeath strains

Glycerol stocks were used to innoculate 2 ml of pH 7 media and grown for approximately 4 hours at 37 °C, until mid-log phase. Cultures were then successively back diluted 100 fold into 2 ml of pH 7 media, and grown for 3 hours. 1-4 steps would be undertaken per day, and at the end of each day glycerol stocks would be made using 1:1 ratio of culture to 50% glycerol. In this way cultures were always kept at either 37 °C or −80 °C and were prevented from ever entering stationary phase and subsequently lowering the pH of the media.

#### Counter and memory strain construction

Actuator, memory and trigger elements were assembled using overlap extension PCR, Gibson assembly, or Golden Gate assembly. Actuator or memory elements were genomically integrated via λ Red recombineering into *E. coli* TB10. From *E. coli* TB10, actuator or memory elements were transduced by P1 transduction into *E. coli* K-12 MG1655. Trigger elements were either integrated into the genome via λ Red recombineering into a TB10 Δ*int/xis* mutant strain, and subsequently transduced by P1 transduction into the K-12 MG1655 strain containing the appropriate actuator or by assembling onto plasmids using a temperature-sensitive, Tn7 transposon insertion backbone derived from pGRG36. From these second group of plasmids, triggers were subsequently integrated into the K-12 MG1655 strain containing the appropriate actuator using the Tn7 transposon. For strain the tc-memory strains: The 434 actuator element was integrated into a TB10 strain via λ Red and subsequently P1 transduced into an *E. coli* NGF-1 strain that contains a λ memory element (Naydich et al., 2019). The trigger element was then integrated into this strain using the Tn7 transposon as described above.

#### Tn7 transposon integration

Trigger plasmids were electroporated into the appropriate actuator-bearing strain. Transformed cultures were grown in SOC media at 30°C for 90 minutes then resuspended in LB with ampicillin and grown at 30°C overnight to select transformants. Cultures were then back-diluted 1:100 into LB containing chloramphenicol and 0.1% arabinose to induce transposase genes while selecting for integration of triggers (which contained a chloramphenicol resistance cassette). After 6 hours at 30°C, cultures were back-diluted 1:100 into LB with chloramphenicol and grown at 42°C for at least 6 hours to cure the temperature-sensitive plasmid while selecting for integration. The back-dilution and curing were repeated a second time. Post-cure cultures were plated on agar containing chloramphenicol, and individual colonies were checked for integration at the attTn7 site by PCR and Sanger sequencing. Plasmid loss was confirmed by restreaming on agar plates containing ampicillin.

#### *In vitro* induction of aTc induced counter and memory strains

For induction time courses, strains were grown in liquid culture for periods of at least 4 hours at each step in the time course. aTc was added or removed from the media. Cultures were back diluted 1:100 between each step, with an extra wash prior to back dilution from +aTc to -aTc cultures. For some replicates, each step was as long as 24 hours. At the end of each step in the time course, 2 µl of each culture was spotted on an agar plate for imaging (with or without inducer and X-gal indicator, as appropriate). For strains with fluorescent reporters, cultures were analyzed by diluting 1:100 in PBS and measuring GFP and mCherry fluorescence using a BD LSR II Flow Cytometer. For strains with a LacZ reporter, cultures were plated on agar plates containing X-gal, and the strain response was assessed by counting blue (on) and white (off) colonies..

#### Imaging

Agar plates were imaged using a custom-built macroscope courtesy of the Kishony Lab and the Harvard Medical School Department of Systems Biology. All spot plates from induction time courses were imaged at 24 hours after plating. Consistent camera settings were used for each channel, and any contrast adjustments were applied consistently to all images in the series.

#### Counter strain trigger excision check by PCR

PCR was used to check for trigger excision as a result of counter induction. For The 434 two-counter with the trigger at the attTn7 locus, the primers used were ADN01 and AND02. For the λ two-counter with the trigger at the arabinose locus, the primers used were ADN03 and AND04. PCR products were run on an agarose gel with a 1 Kb Plus Ladder (ThermoFisher #10787018).

#### Construction of tc-P*_asr_*AT09:GFP

Trigger element was constructed using a golden gate reaction of the products of three PCR reactions. Primer TC36 and TC37 using acidTRP-09 genomic DNA as a template, TC38 and TC39 using tc-P*_tet_*:mCherry genomic DNA as a template, TC40 and TC47 using plasmid pFS01 (submitted to Addgene) as a template. The resultant plasmid was transformed into an *E. coli* K12 strain with the 434 actuator element integrated into the genome, resulting in strain tc-P*_asr_*AT09:GFP.

#### Construction of tc-acidTRP09

The sequence in Table S4 was ordered as a gblock from Integrated DNA Technologies. This dsDNA was then transformed into TB10 using previously stated protocols, and the relevant region was transferred to into an *E. coli* K12 strain with the 434 actuator element integrated into the genome via P1 transduction, resulting in strain tc-acidTRP-09.

#### pH time course assay for tc-P*_asr_*AT09:GFP two counter constructs

tc-P*_asr_*AT09:GFP and P*_asr_*AT09:GFP glycerol stocks were used to inoculate pH 7 media and grown overnight at room temperature to late log phases. Cultures were then back diluted 1000 fold into either pH 7 or pH 5 media, and grown for 5 hours at 37 °C. cultures were again back diluted 1000 fold into pH 7 or pH 5 media and grown for 5 hours at 37 °C. A final 1 to 1000 back dilution into pH 7 or pH 5 media and 5 hour growth step at 37 °C was conducted. After each step a glycerol stock was made by combining cultures with 50% glycerol in a 1:1 ratio. Glycerol stocks were then defrosted and diluted 100 fold and assayed on a flow cytometer.

#### pH time course assay for tc-P*_asr_*AT09-memory

tc-P*_asr_*AT09-memory glycerol stocks were used to inoculate pH 7 media and grown overnight at room temperature to late log phase. four subsequent 5 hour growth steps at 37 °C were conducted, each one a successive 1000 fold dilution into alternating pH 5 then pH 7 media. Glycerol stocks were made after a growth step if it could not all be carried out on the same working day. After each growth step, a dilution series in the relevant media was made and plated onto pH 7 buffered plates supplemented with X-gal. the percentage of blue colonies was calculated and used to determine the percentage of cells expressing LacZ.

#### pH time course assay for tc-acidTRP-09

tc-acidTRP-09 glycerol stocks were used to inoculate pH 7 media and grown overnight at room temperature to late log phase. To assay the survival ratio after successive growth steps, overnight culture was back diluted 1000 fold into pH 5 media and grown for 5-10 hours at 37 °C. this was then back diluted 1000 fold into pH 7 media and grown for 5 hours at 37 °C, and then again into pH 5 media for a final 5 hour growth step at 37 °C. Glycerol stocks were made after a growth step if it could not all be carried out on the same working day. After each growth step, a dilution series in the relevant buffered media was made and plated on pH 7 and pH 5 plates as described for survival assays. To assay for survival ratio after varying sequences of growth conditions, the overnight culture was back diluted into both pH 7 or pH 5 media and grown for 5 hours at 37 °C for two more consecutive growth steps, resulting in four different sequences of conditions. Each of these four cultures was plated on both pH 7 and pH 5 plates, and a survival ratio was calculated for each.

## Supplementary Figures

**Figure S1.**
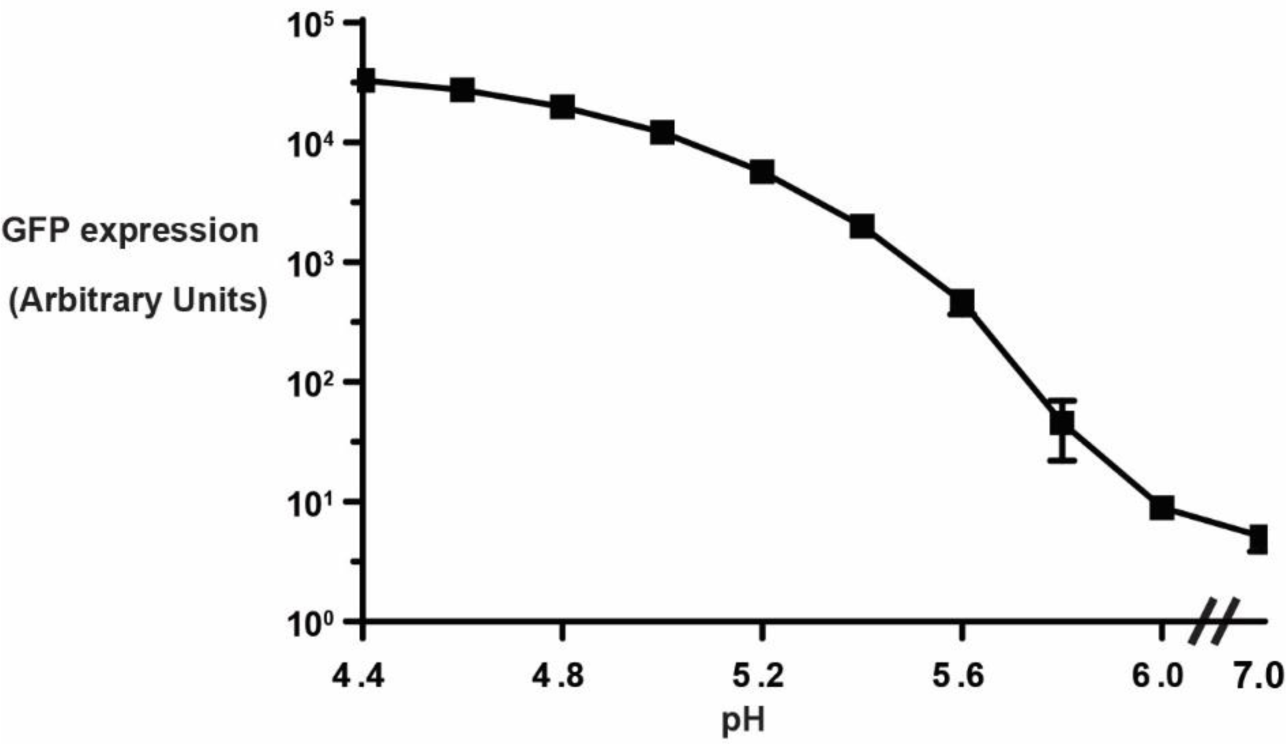
Related to Figure 1. Induction curve of P*_asr_*AT09:GFP in M9 media with 1 g/L yeast extract buffered to a range of pH’s. GFP fluorescence was measured using a flow cytometer after five hours of growth at 37 °C. data points are from three technical repeats, error bars are standard deviation.

**Figure S3.**
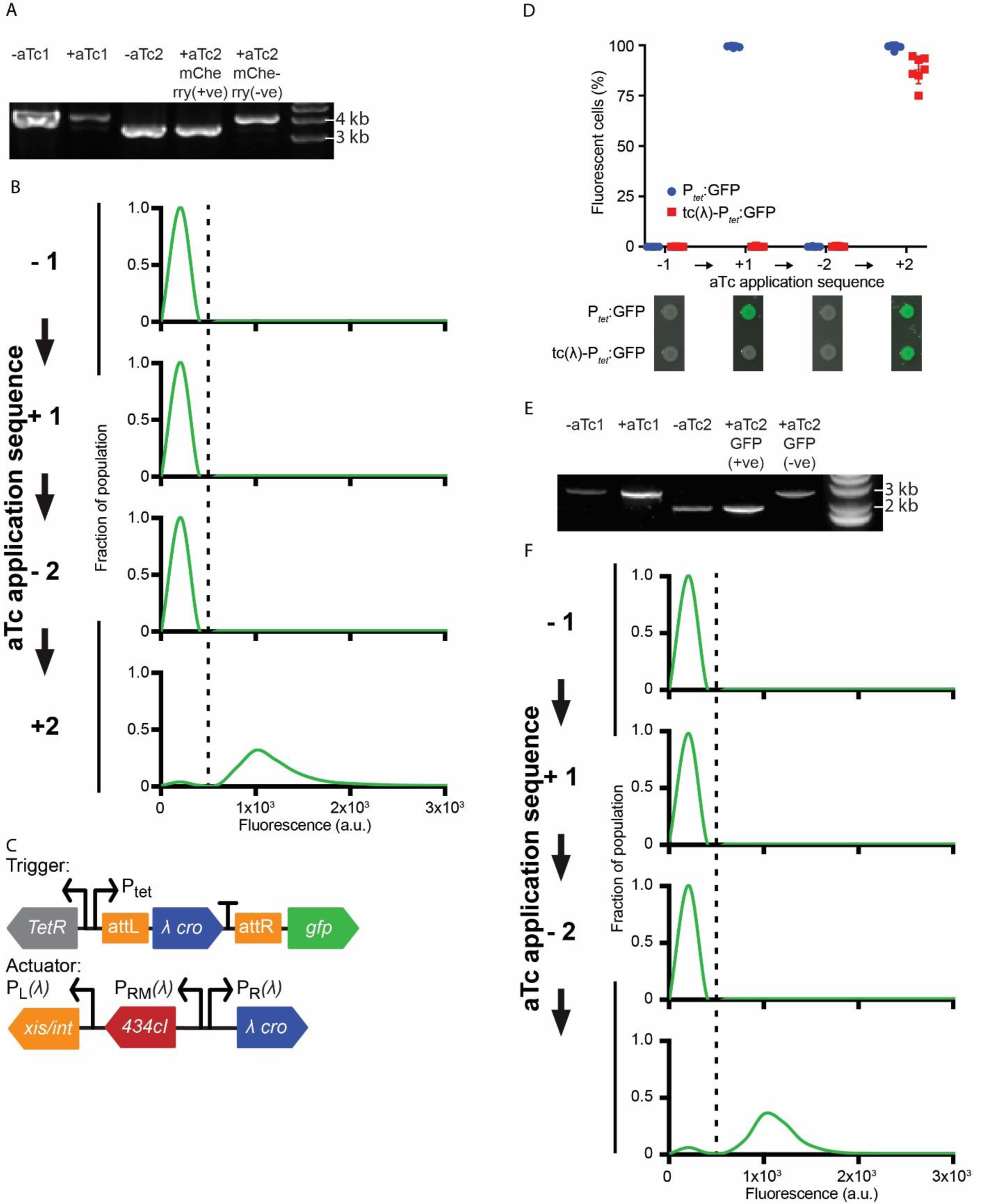
Related to Figure 3. S3A) Agarose gel depicting bands from PCR amplification of the 434 two-counter trigger during a time course of aTc induction. At each step of the time course, genomic DNA from a colony of tc-P*_tet_*:mCherry was used as a template for PCR at the Tn7 locus to check for excision of the trigger. (Full trigger length: 4.0 kb; excised trigger length: 3.3 kb.) Rightmost band corresponds to a single mCherry-negative colony picked from the +aTc2 plate. S3B) Distribution of mCherry fluorescence intensity in a single culture of the tc-P*_tet_*:mCherry, in which 97% of the population has successfully undergone trigger exicision as a result of the first aTc stimulus. The cutoff between mCherry negative and mCherry positive cells is indicated by the dotted line at ∼100 a.u. S3C) λ two-counter circuit with a GFP reporter (tc(λ)-P*tetR*:GFP). S3D) Response of tc(λ)-P*_tet_*:GFP to an aTc induction time course with two stimulus pulses. aTc was applied and washed out for growth periods of at least 4 hours (+1 and +2 growth steps). A P*_tet_*:GFP strain is shown as a control. Error bars represent SD of seven biological replicates. Spots of cultures on agar media (with or without aTc) are shown below at each step of the time course. Spot images consist of a GFP color overlay on a grayscale brightfield image. S3E) Agarose gel depicting bands from PCR amplification of tc(λ)-P*_tet_*:GFP at the araB/araC locus. At each step of the time course, genomic DNA from a colony of tc(λ)-P*_tet_*:GFP was used as a template for PCR to check for excision of the trigger. (Full trigger length: 2.5 kb; excised trigger length: 1.8 kb.) Rightmost band corresponds to a single GFP-negative colony picked from the +aTc2 plate. S3F) Distribution of GFP fluorescence intensity in a single culture of the tc(λ)-P*_tet_*:GFP, in which 93% of the population has successfully undergone trigger excision as a result of the first aTc stimulus. The cutoff between GFP- and GFP+ cells is indicated by the dotted line at ∼100 a.u.

**Figure S4.**
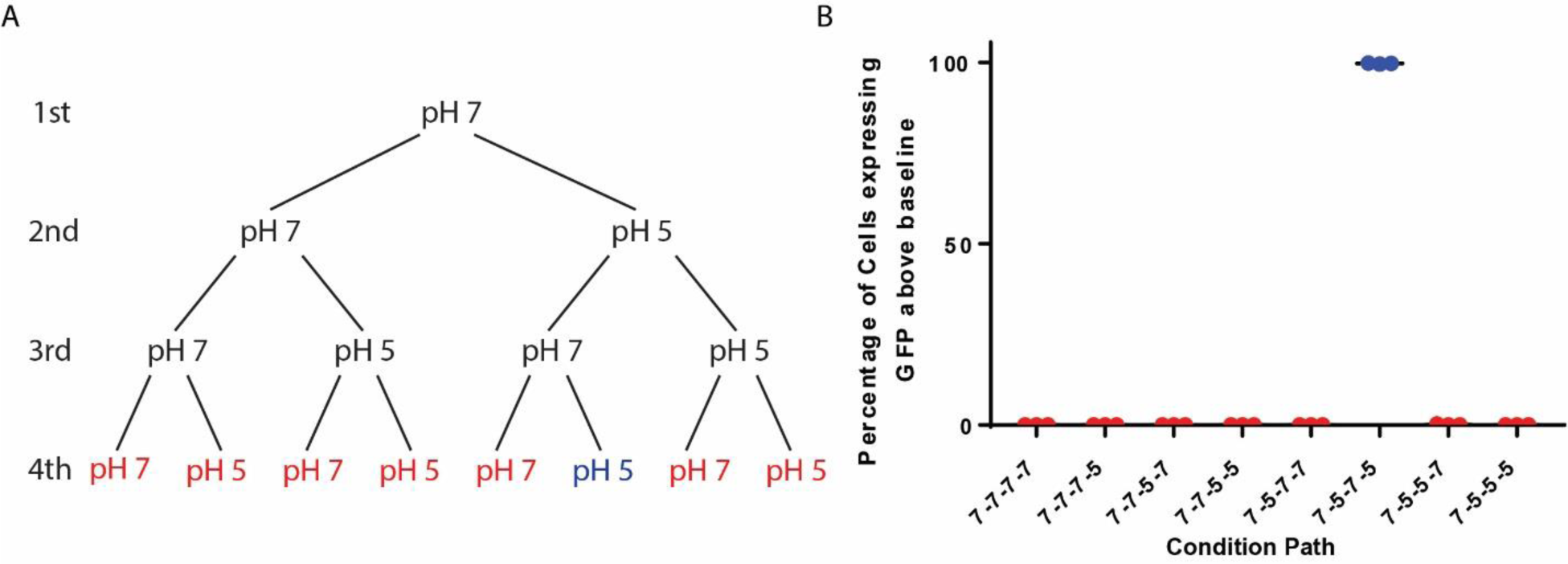
**A)** Tree of conditions for acid sensitive two count assay. each level of the tree represents a growth step, with each step after the first having one growth in pH 7 conditions and one in pH 5. On the final level, only the condition highlighted in blue would expect to express the reporter gene. All other conditions (red) would expect no expression. **B)** percentage of cells expressing GFP above that of a negative controls (wild type and uninduced P*_asr_*:GFP). Only cells exposed to two non-consecutive growth steps in pH 5 conditions showed a significant proportion of the population expressing GFP.

**Table S1. related to Figure 1.**
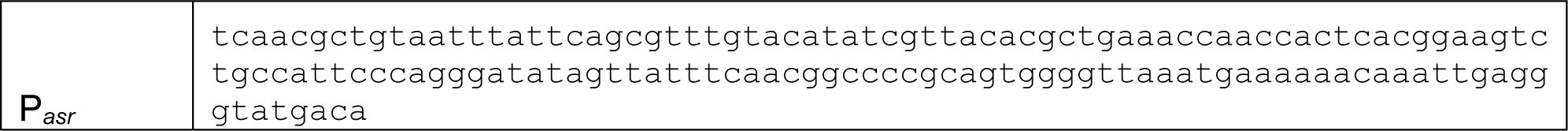
Sequence of promoter and ribosome binding site of P*_asr_* ordered from SGI DNA.

**Table S2, related to Figure 1.**
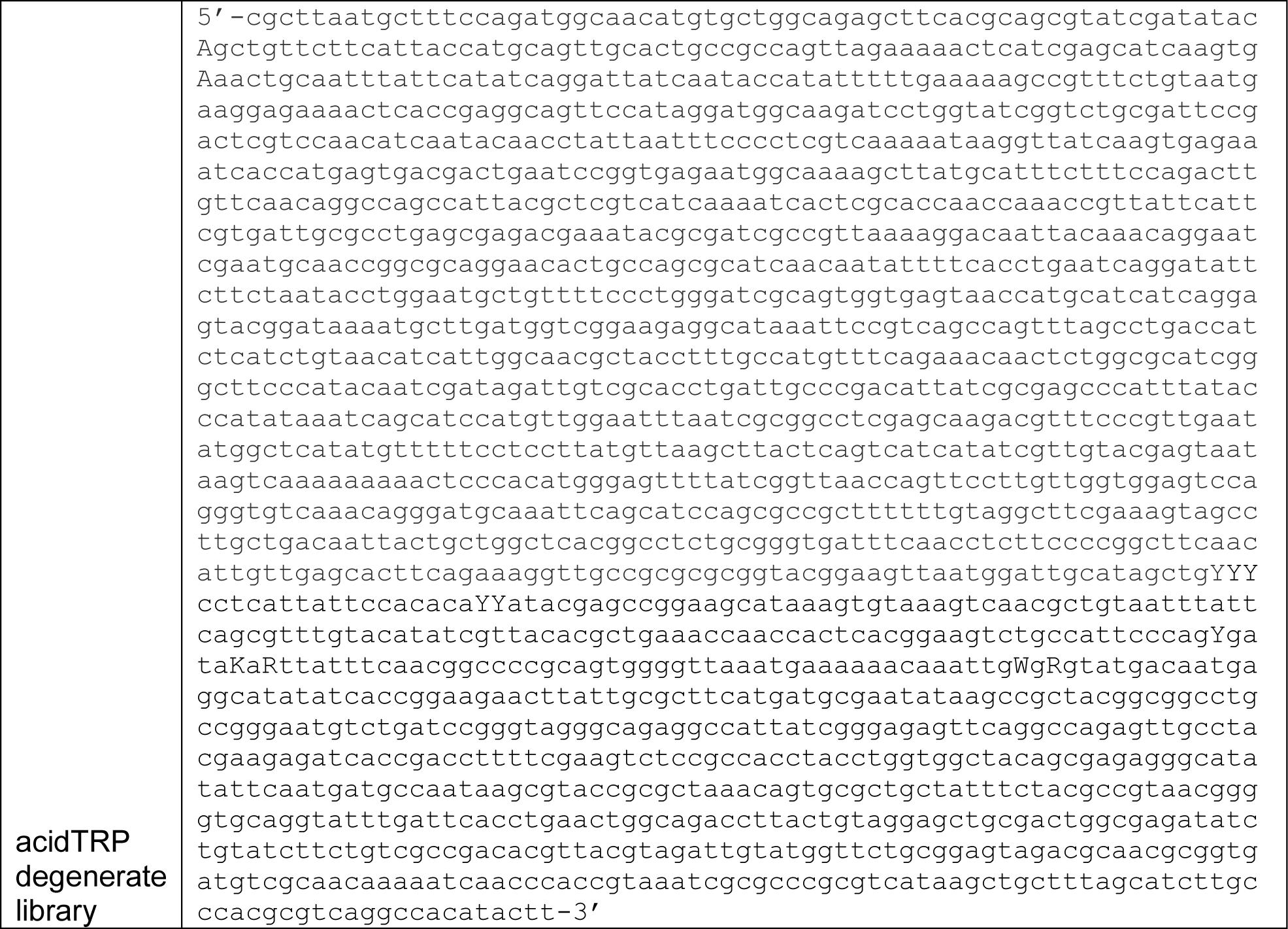
Full sequence of degenerate acidTRP library ordered from SGI DNA. Degenerate bases are capitalised.

**Table S3. Related to Figure 1.** Identity of degenerate bases for acidTRP candidates reported on in Figure 1C. addition modifications are described under notes, with the bp position referencing the sequence in Table S2.

|  | Y | Y | Y | Y | Y | Y | K | R | W | R | Notes |
| --- | --- | --- | --- | --- | --- | --- | --- | --- | --- | --- | --- |
| acidTRP-TM | T | T | T | T | C | C | G | G | A | A |  |
| acidTRP-01 | T | C | C | C | C | C | G | G | A | G |  |
| acidTRP-02 | T | T | T | C | C | T | T | G | A | A |  |
| acidTRP-03 | C | T | T | C | T | C | G | G | A | A |  |
| acidTRP-04 | T | C | C | C | C | T | G | G | A | A |  |
| acidTRP-05 | C | T | C | T | T | T | G | A | A | G | A 1278 deleted |
| acidTRP-06 | C | T | C | T | C | T | T | G | A | G |  |
| acidTRP-07 | T | C | C | C | C | T | T | A | A | G | T 1346 deleted |
| acidTRP-08 | T | T | C | C | C | C | T | G | T | A |  |
| acidTRP-09 | T | T | C | T | C | T | G | G | A | G |  |
| acidTRP-10 | T | T | C | C | C | C | T | A | A | A |  |
| acidTRP-11 | T | T | C | C | T | C | T | G | T | G |  |
| acidTRP-12 | T | T | T | C | C | T | G | A | A | A |  |
| acidTRP-13 | T | T | T | C | C | T | T | A | A | A |  |

**Table S4. Related to Figure 5.**
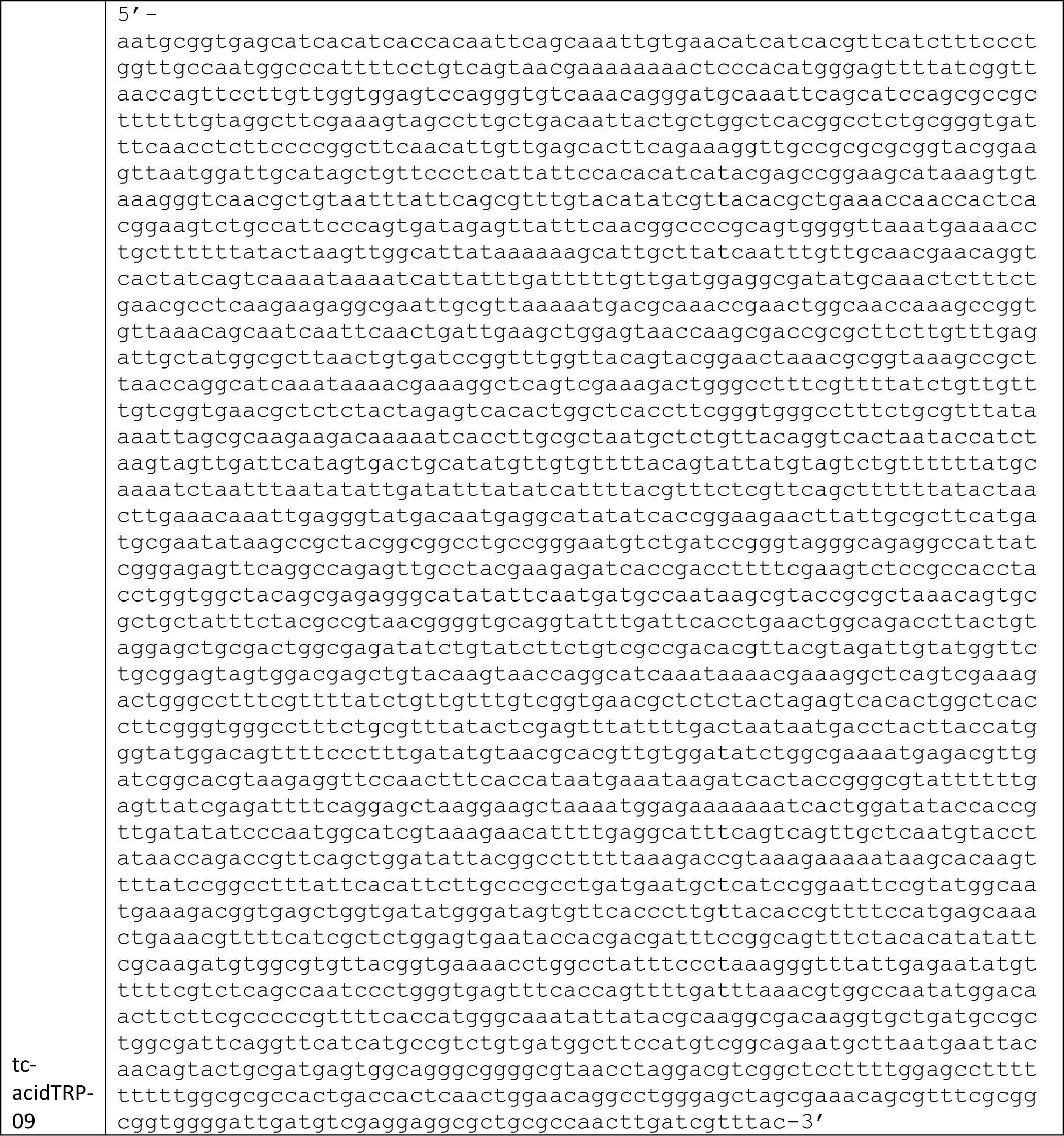
Sequence of tc-acidTRP-09 ordered as gblock from lOT.

**Table S5. Related to Methods.**
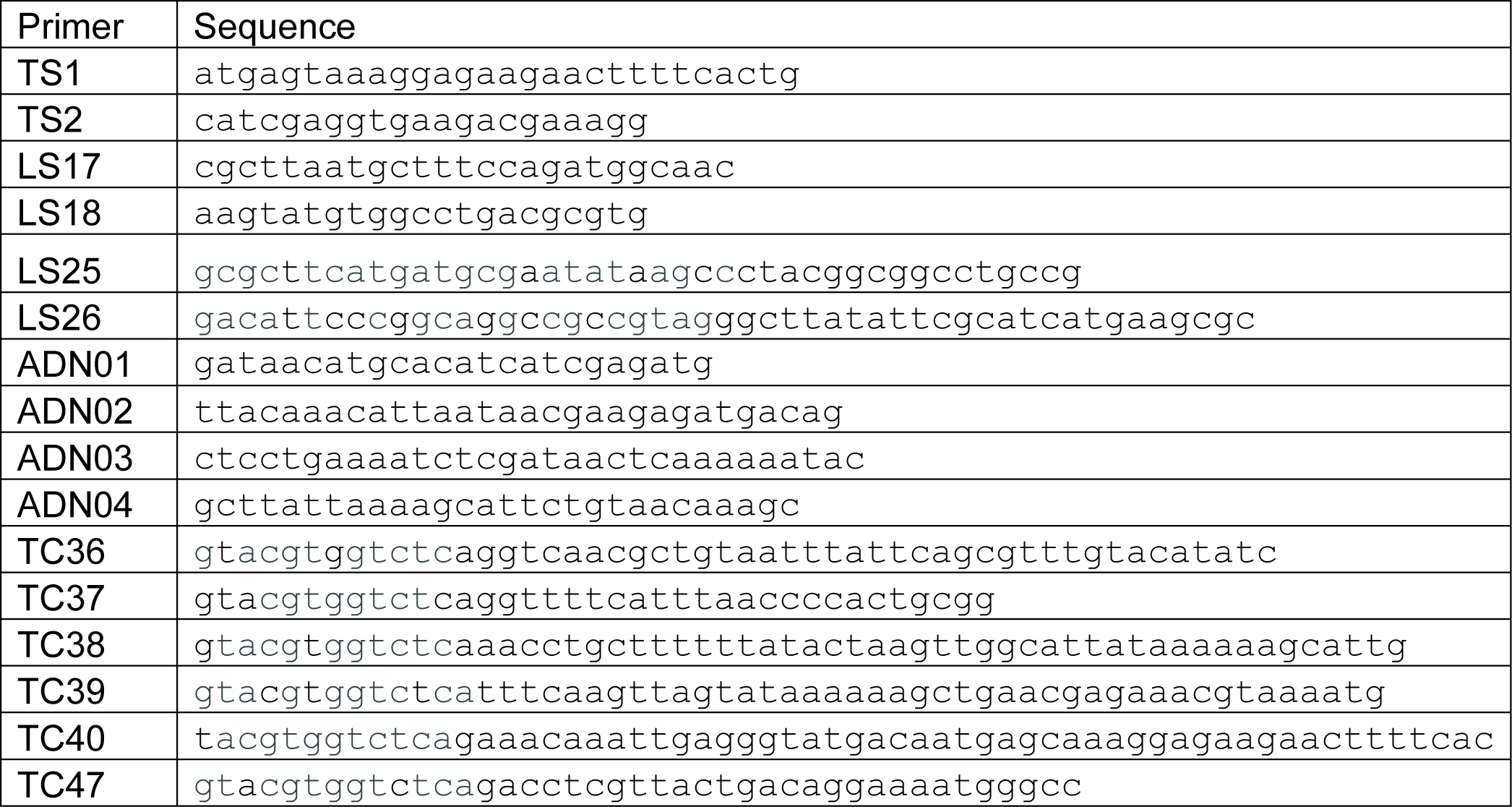
Full sequence of all oligos described in the methods.

